# *In Silico* Investigations on Curcuminoids from *Curcuma longa* as Positive Regulators of Wnt/β-catenin Signaling Pathway in Wound Healing

**DOI:** 10.1101/2020.03.19.998286

**Authors:** Riyan Al Islam Reshad, Sayka Alam, Humaira Binte Raihan, Kamrun Nahar Meem, Fatima Rahman, Fardin Zahid, Md. Ikram Rafid, S. M. Obaydur Rahman, Sadman Omit, Md. Hazrat Ali

## Abstract

*Curcuma longa* (Turmeric) is a traditionally used herb in wound healing. The efficacy of fresh Turmeric paste to heal wound has already been investigated in multiple ethnobotanical studies. Wnt/β-catenin signaling pathway plays a significant role in wound healing and injury repair processes which has been evident in different *in vitro* studies. This study aims to analyze the potentiality of Curcuminoids (Curcumin I, Curcumin II and Curcumin III) from *Curcuma longa* to bind and enhance the activity of two intracellular signaling proteins-Casein Kinase-1 (CK1) and Glycogen Synthase Kinase-3β (GSK3B) involved in Wnt/β-catenin signaling pathway. Present study is largely based on computer-based molecular docking program which mimics the *in vivo* condition and works on specific algorithm to interpret the binding affinity and poses of a ligand molecule to a receptor. Curcumin I showed better affinity of binding with CK1 (−10.31 Kcal/mol binding energy) and Curcumin II showed better binding affinity (−7.55 Kcal/mol binding energy) for GSK3B. Subsequently, Drug likeness property, ADME/Toxicity profile, Pharmacological activity and Site of metabolism of the Curcuminoids were also analyzed. All of the ligand molecules showed quite similar pharmacological properties.

## 1. Introduction

### 1.1. Wound, Wound Healing Process and Treatments

Skin is the largest organ of the body which act as a primary protective barrier. Any physical damage on the skin, causes exposure of subcutaneous tissue following a loss of skin integrity known as wound. Wound provides a moist, warm, and nutritious environment that is suitable for microbial colonization and proliferation [1]. Infected wounds harbor diverse populations of microorganisms. However, in some cases these microorganisms can be difficult to identify and fail to respond to antibiotic treatment, resulting in chronic non-healing wounds due to the formation biofilm [2].

Wound healing takes place by three main events: Inflammatory phase, Proliferative phase and Maturation and remodeling phage. During the inflammatory phase there is an influx of inflammatory cells and local Wnt/β-catenin signaling begins to increase [3]. Wnt proteins are glycoproteins that regulate cell proliferation, migration and specification of cell fate. Wnt proteins are classified according to their ability to promote stabilization of β-catenin in the cytoplasm. The β-catenin-dependent Wnt pathway signals through cytoplasmic stabilization and accumulation of β-catenin in the nucleus to activate gene transcription which leads to cell division and cell proliferation [4]. During the proliferative phase a scab (eschar) is formed and the wound is reepithelialized. This phase includes an increased local Wnt response, which increases the accumulation of β -catenin in the cytoplasm. Increases in β -catenin level leads to some gene transcriptions, such as the matrix metalloproteinases, which causes extracellular matrix deposition, angiogenesis, and the recruitment and proliferation of multiple cell types including stem cells, keratinocytes, and fibroblasts. The third phase, the maturation and remodeling phase, is characterized by extensive extracellular matrix remodeling [3][5].

Lots of treatments for wound care are available in the market. One of most commonly used is Standard wound healing technique and wound dressing. A wound dressing includes a cover membrane comprising a semi-permeable material with an adhesive-coated skin contact surface. An intermediate layer of material may be placed between the wound and the membrane contact surface for either absorbing fluids from the wound, e.g. with a hydrocolloid or hydrophilic material, or for passing such fluids to the opening with a synthetic material, e.g. rayon [6]. On other hand standard wound technique includes wet-to-dry dressings, gel and sliver sulfadiazine (Slivadene). Another method is Vacuum Assisted Closure (VAC) therapy, which is most effective than standard wound healing. This therapy accelerates the healing process by removal of edema and stimulation of mitosis through cell deformation. Success of the treatment depends on the condition of the wound [7]. 1% silver sulfadiazine, an antimicrobial topical ointment, is one the popular medicine in treatment of burn wounds around the world. It is easy and convenient to use, causes no pain, low toxicity and sensitivity. However, studies have shown that this drug also causes side effects such as reduction of white blood cells, toxic epidermal necrolysis, increased skin pigmentation, neutropenia, increased bacterial resistance and the skin appearance would not return to normal. For some countries, like Iran raw materials are often needed to be imported at higher prices [8]. Moreover, an estimated US$ 25 billion is spent annually only in USA for the wound healing purposes and the margin is increasing day by day due to the increasing healthcare cost. While current therapeutic agents have lower efficacy and many adverse side effects, medicinal plants which are being used as medicine from very ancient time have been proven effective and safe for wound healing and thus herbal medicine can be a lucrative alternative of current therapeutic agents [9]. Turmeric is extensively used as spice, preservatives and coloring material in food. Numerous studies have been conducted with Turmeric over the last few decades which have proven its multiple functions in combating multiple diseases. It contains Sabinene, Borneol, Zingiberene and some other major phytochemicals of great therapeutic values as well as Curcuminoid which are responsible for the yellow color of Turmeric and it comprises Curcumin I, II and III [10] [11]. Role of both Turmeric and individual Curcumin from the plant in wound healing has already been demonstrated in laboratory experiments [10][12].

### 1.2. Wnt/β-catenin Pathway and Its Involvement in Wound Healing

The Wnt/β-catenin signaling pathway is activated by the binding of secreted Wnt proteins to a receptor complex containing a member of the FZD family and LRP5/6 [13]. A number of complex proteins facilitate the Wnt/β-catenin Pathway. Casein Kinase-1 (CK1) and Glycogen Synthase Kinase-3β (GSK3B) phosphorylate several important components like B-catenin, Axin and APC (Adenomatous Polyposis Coli), in the Wnt/β-catenin signaling pathway and act as either negative or positive regulators of the pathway. These phosphorylation events result in tighter association of Axin and APC with b-catenin CK1 is a family of serine/threonine-specific protein kinases regulates diverse cellular processes like Wnt signaling [14]. They phosphorylate several pathway components and exert a dual function, acting as both Wnt activators and Wnt inhibitors. The CK1 family consists of six human isoforms (α, δ, ε, γ1, γ2, γ3) are ubiquitously expressed and share a highly homologous kinase domain, flanked by variable N-terminal and C-terminal extensions. The amino-terminal kinase domain is highly conserved in all CK1 family members [13] [15]. The serine/threonine kinase GSK3B binds to and phosphorylates several proteins in the Wnt pathway and is serving to the down regulation of β-catenin. As a negative regulator of Wnt signaling, GSK3B would qualify as a potential tumor suppressor [16].

#### 1.2.1. Wnt/β-catenin Pathway in Absence of Wnt Ligand

In absence of Wnt ligand, β-catenin is phosphorylated and targeted to degradation by a protein complex consisting of several molecules including Axin, APC, CK1, and GSK3B. Axin is a protein which serves as a scaffold for formation of destruction complex (**Figure 1**) [17]. APC is a large protein that interacts with both β-catenin and Axin. It contains three Axin-binding motifs that are interspersed between a series of 15 and 20 amino acid repeats[18].The scaffolding protein Axin and tumor suppressor APC form a β-catenin destruction complex that binds cytosolic β-catenin and sequentially phosphorylate specific N-terminal residues of β-catenin by serine/threonine kinases, casein kinase 1γ (CK1γ) and glycogen synthase kinase 3β (GSK3B) [19] [20]. Once β-catenin is hold into the destruction complex by APC and phosphorylated by GSK3B, the phosphorylated β -catenin is a target for β-Trcp which is E3-ubiquitin ligase. β-Trcp transfers ubiquitin chain to the β-catenin and ubiquitinated β-catenin degrades via ubiquitin proteasome mediated pathway [18] [21]. In the nucleus, the Wnt target genes are bound by transcription factors of TCF/ LEF. In absence of β-catenin these TCF/LEF transcription factors are bound to Groucho which is a transcriptional repressor, does not allow the Wnt target genes to express [22][23].

**Figure 1:**
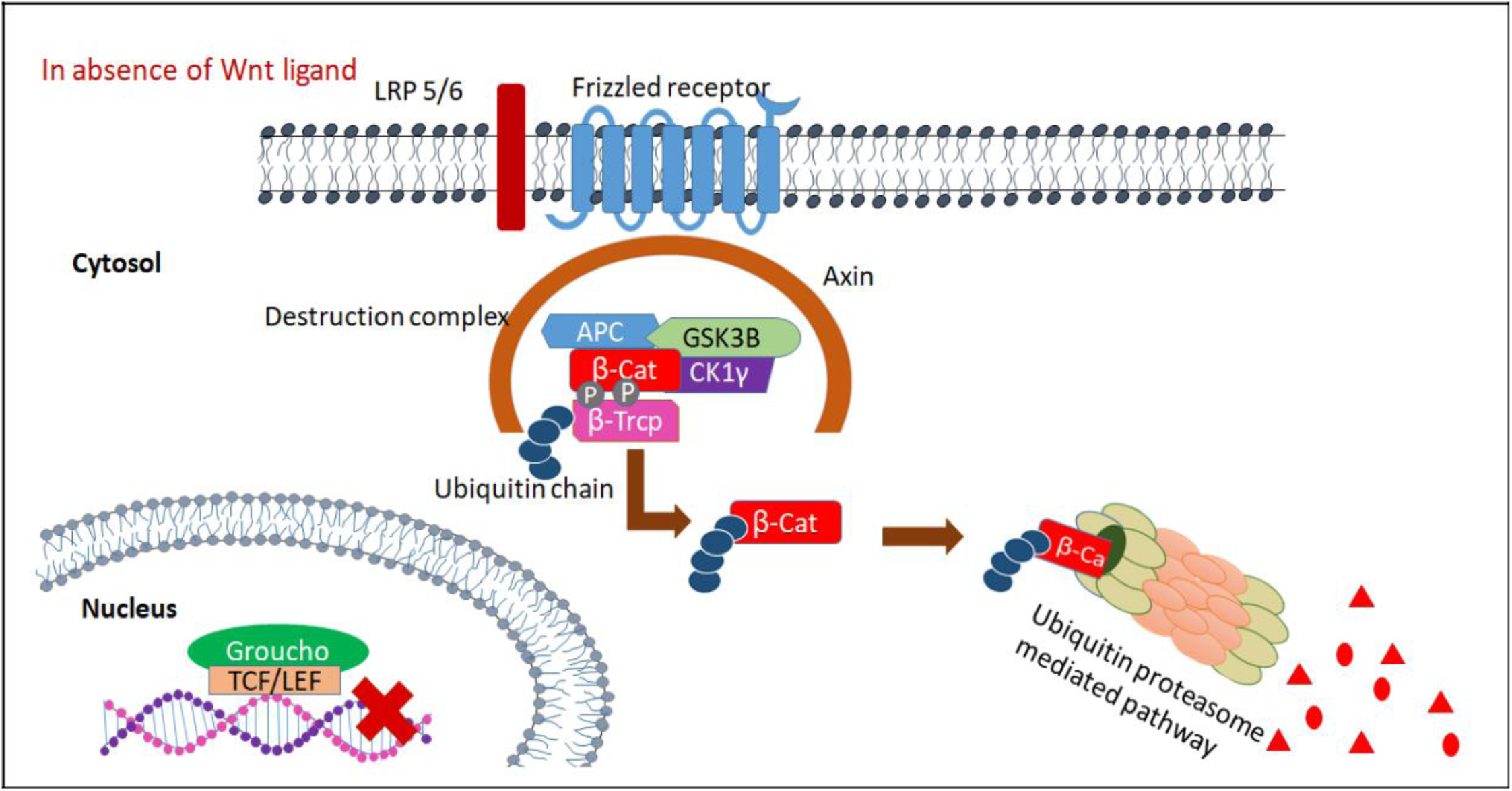
In the absence of Wnt proteins, Axin, APC, β-catenin, **and** GSK3B form a complex, in which β-catenin is phosphorylated by CK1 and GSK3B, leading to β-catenin degradation via Ubiquitin proteasome mediated pathway.

#### 1.2.2. Wnt/β-catenin Pathway in Presence of Wnt Ligand

In presence of Wnt ligand, The Wnt signaling cascade is triggered upon binding of members of the Wnt family proteins to a co-receptor complex, including frizzled (Fz, a G protein-coupled receptor-like protein) and LRP5 or -6 (**Figure 2**). The signal is transmitted through recruitment of several proteins to the C-terminal intracellular moieties of the activated Fz and LRP5/6 coreceptors. From this point, binding Wnt ligand to the frizzled receptor leads to dimerization of LRP 5/6 with frizzled receptor. Dishevelled (DVL), a cytoplasmic binding protein which binds to the frizzled receptor, leads to instability of destruction complex [24]. Disheveled (Dvl) is recruited and posttranslationally modified, and depending on the specific nature of the Wnt and of the Fz that are complexed with LRP5/6, three independent pathways can be activated: canonical, noncanonical, or Ca2 [25].This instable destruction complex cannot hold β-catenin any longer. As APC cannot hold β-catenin, β-catenin is no longer phosphorylated by GSK3B. And B-Trcp protein cannot add ubiquitin chain to β-catenin. As a result, β-catenin is degraded by Ubiquitin proteasome mediated pathway. Thus β-catenin is released to the cytosol and amount of β-catenin in the cytosol increases [23] [26]. β-catenin migrates to the nucleus and replace the Groucho repressor. The interaction of β-catenin with the N terminus of Tcf converts it into an activator, translating the Wnt signal into the transient transcription of Tcf target genes [27]. β-catenin also recruits CBP and Brg1. These are transcriptional activator which actives the transcription of Wnt-ligand targeted genes. These Wnt-targeted genes helps in cell cycle progression from G1 phase to S phase, which leads to cell division and cell proliferation of stem cells, keratinocytes and fibroblasts. The activation of the Wnt pathway has a key role for fibroblast activation and collagen release in fibrosis. Wnt signaling stimulated the differentiation of resting fibroblasts into myofibroblasts, increased the release of extracellular matrix components and induced fibrosis. Stem cells, Fibroblasts and keratinocytes migrates to the wounded site as immune response. As a result, scar formation occurs at the wounded site and injury repairs [3] [21] [28].

**Figure 2:**
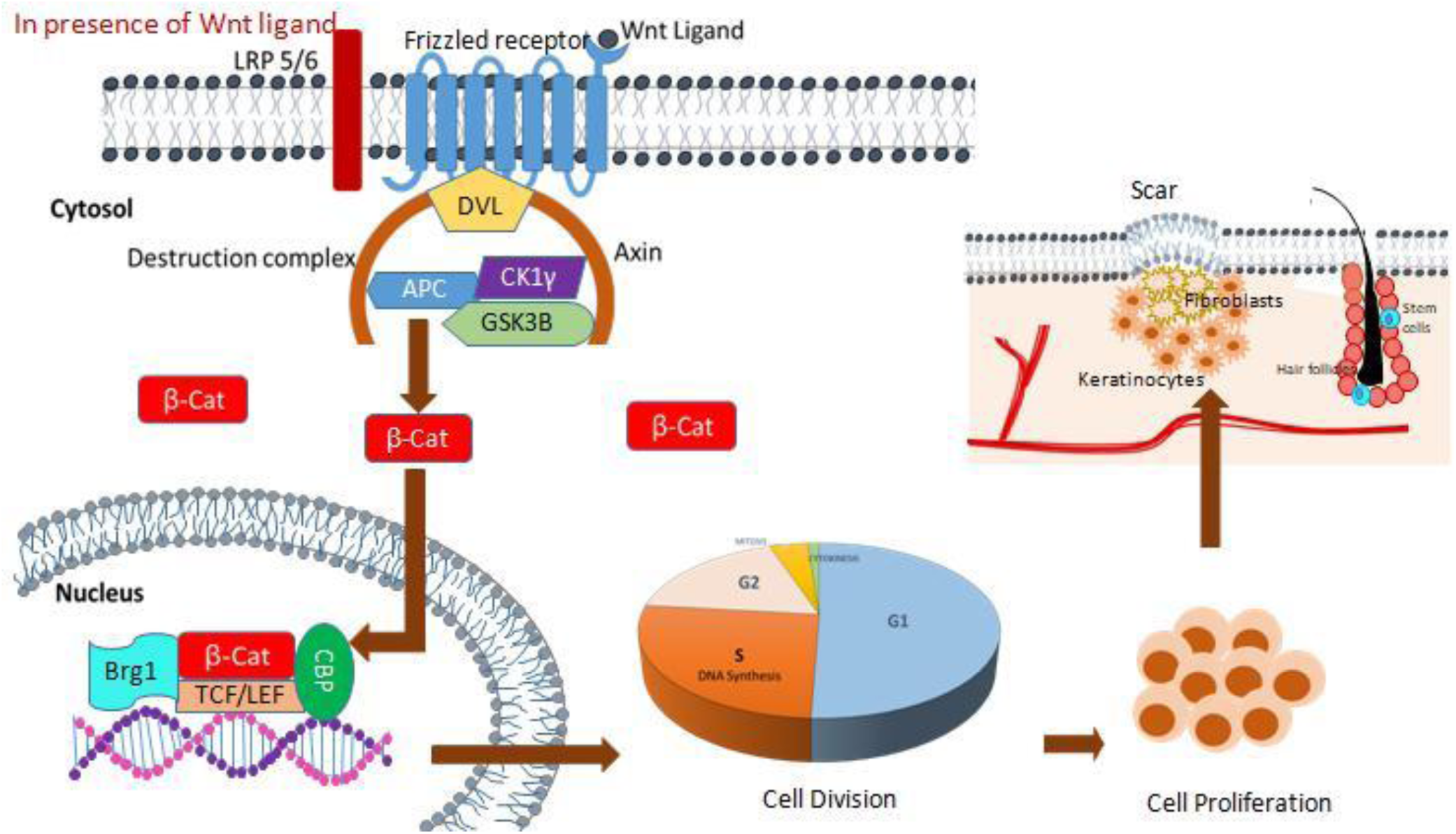
Wnt proteins bind to Frizzled (Fz) and LRP-5, Axin is recruited to the membranes. The interaction between Axin and LRP-5 may prevent Axin from participating in the degradation of β-catenin. Dvl may also receive signals from the Fz/LRP complex, resulting in the inhibition of GSK3B. β-catenin is not degraded, which eventually leads to transcription of the Wnt-targeted gene, which results in following event like cell division, cell proliferation and cell migration to the wound site.

In this experiment, Curcumin I, II and III (**Figure 3**) have been docked with two intended targets of Wnt/ β-catenin signaling pathway-CK1 and GSK3B (**Figure 4**) based on the hypothesis that, these phytocompounds might bind to the targets and act as a positive regulators of those targets which may lead to increased signaling and eventually effective wound healing process. Later on, druglikeness property, ADME/Toxicity test, Pharmacological activity and Site of metabolism of the selected phytocompounds were also analyzed.

**Figure 3:**
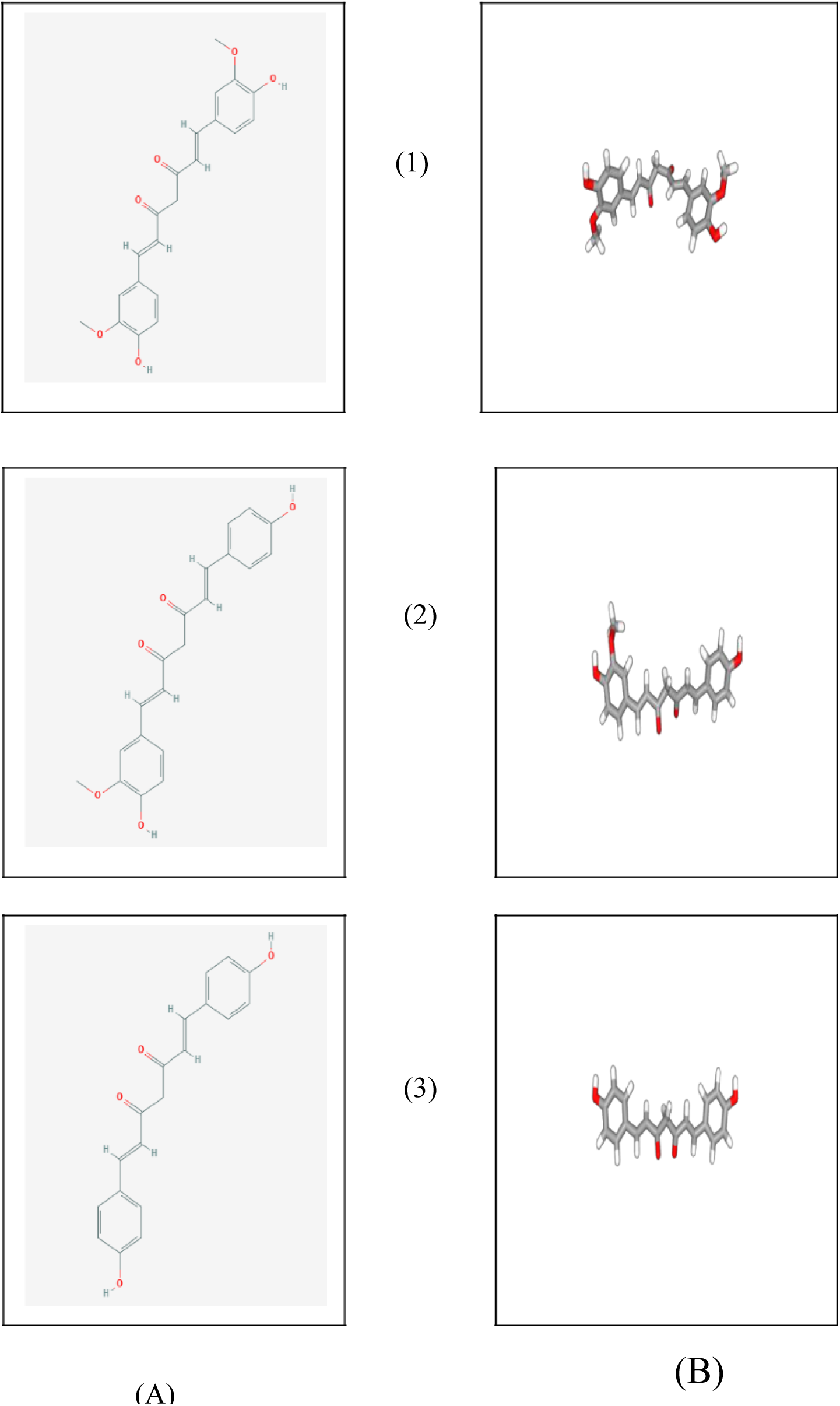
Structures of (1) Curcumin I (PubChem CID: 969516), (2) Curcumin II (PubChem CID: 5469424) and (3) Curcumin III (PubChem CID: 5315472); (A) 2D, (B) 3D.

**Figure 4:**
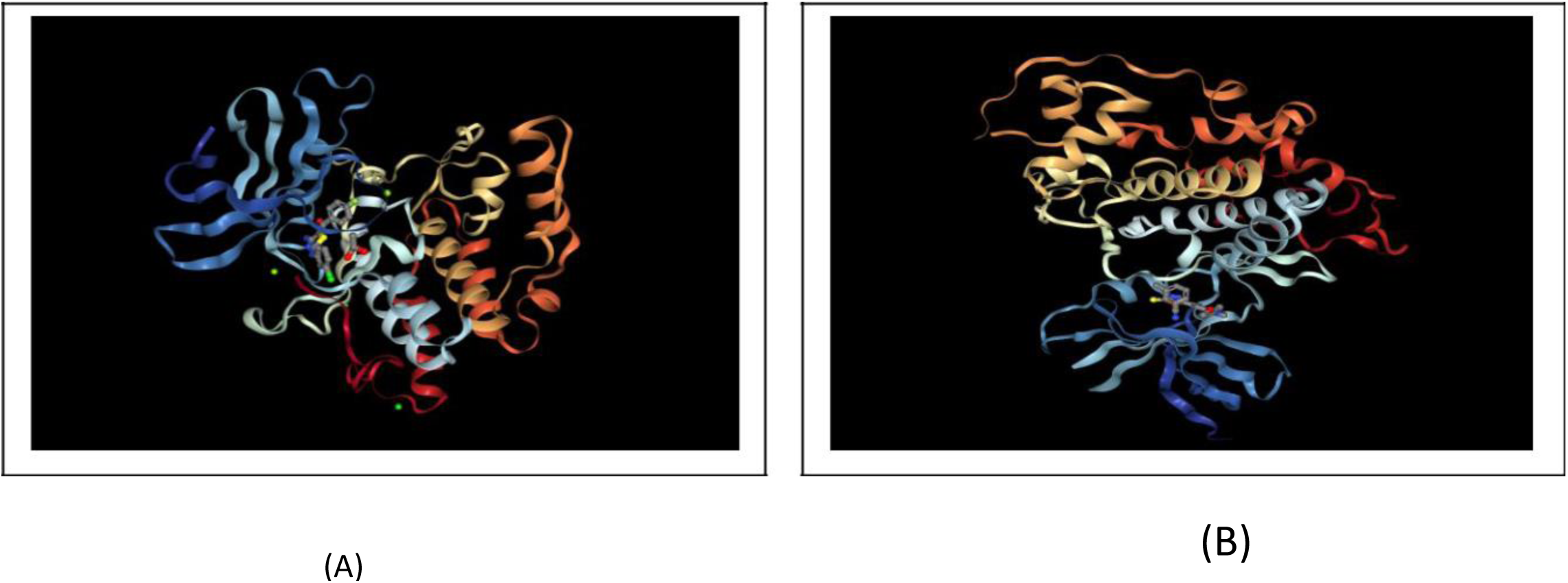
Three dimension structure of (A) Casein Kinase-1 (PDB Id: 2IZS), (B) Glycogen Synthase Kinase-3β (PDB Id: 3F88) in their native ligand bound form. Receptors are represented in cartoon style and ligands are represented in ball and stick style.

## 2. Materials and Methods

### 2.1. Molecular Docking

#### 2.1.1. Protein Preparation

Three dimentional structures of Casein Kinase 1 gamma (PDB Id: 2IZS) and Glycogen Synthase Kinase-3 Beta (PDB Id: 3F88) were downloaded in PDB format from Protein Data Bank (www.rcsb.org). The structures were then prepared and processed using the Protein Preparation Wizard in Maestro Schrödinger Suite (v11.4) [29]. Bond orders were assigned to the structures, hydrogens were added to heavy atoms. All of the water molecules were erased from the atoms and selenomethionines were changed over to methionines. At last, the structures were refined and after that minimized utilizing default Optimized Potentials for Liquid Simulations force field (OPLS_2005). Minimization was performed setting the greatest substantial particle RMSD (root-mean-square-deviation) to 30 Å and any extraordinary water under 3H-bonds to non water was again eradicated during the minimization step.

#### 2.1.2. Ligand Preparation

The 3D conformations of Curcumin I (PubChem CID: 969516), Curcumin II (PubChem CID: 5469424) and Curcumin III (PubChem CID: 5315472) were downloaded from PubChem (www.pubchem.ncbi.nlm.nih.gov). These structures were then processed prepared using the LigPrep wizard of Maestro Schrödinger suite [30]. Minimized 3D structures of ligands were generated using Epik2.2 and within pH 7.0 +/- 2.0 in the suite. Minimization was again carried out using OPLS_2005 force field which generated maximum 32 possible stereoisomers depending on available chiral centers on each molecule.

#### 2.1.3. Receptor Grid Generation

Grid usually restricts the active site to specific area of the receptor protein for the ligand to dock specifically within that area. In Glide, a grid was generated using default Van der Waals radius scaling factor 1.0 and charge cutoff 0.25 which was then subjected to Optimized Potentials OPLS_2005 force field for the minimized structure. A cubic box was generated around the active site (reference ligand active site) of target molecules. Then the grid box dimension was adjusted to 10 Å ×10 Å ×10 Å for docking to be carried out.

#### 2.1.4 Glide Standard Precision (SP) Ligand Docking

SP adaptable glide docking was carried out using Glide in Maestro Schrödinger [31]. The Van der Waals radius scaling factor and charge cutoff were set to 0.80 and 0.15 respectively for all the ligand molecules under study. Final score was assigned according to the pose of docked ligand within the active site of the receptor molecules. The docking result is summarized in **Table 1**. Best possible poses and types of ligand-receptor interactions (**Figure 5 and 6**) were analyzed utilizing Discovery Studio Visualizer [32].

**Table 1:**
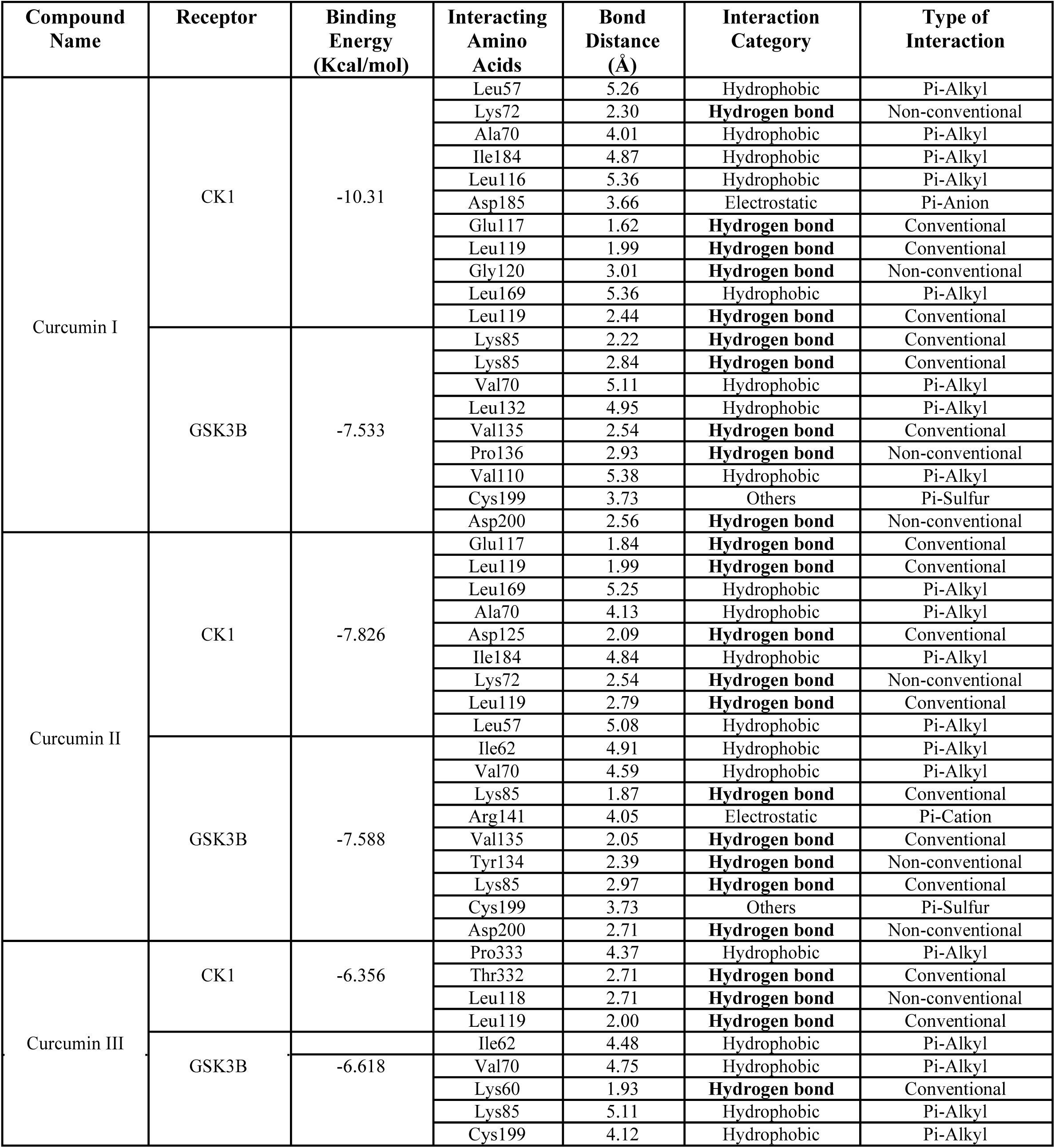
Result of molecular docking between Curcumin I (PubChem CID: 969516), Curcumin II (PubChem CID: 5469424) Curcumin, III (PubChem CID: 5315472) and CK-1 (PDB Id: 2IZS) and GSK3B (PDB Id: 3F88).

**Figure 5:**
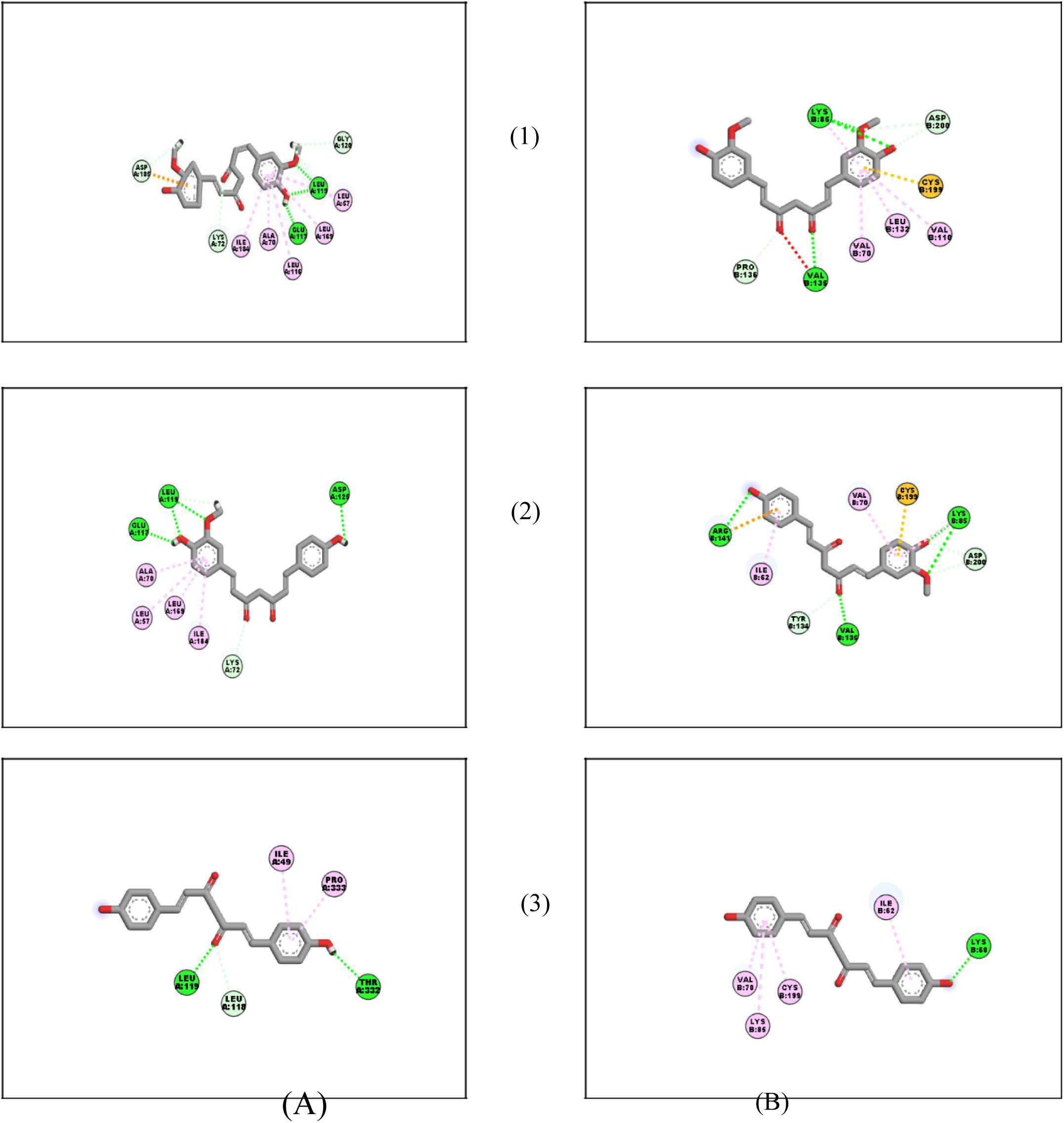
2D representation of ligand-receptor interactions between (1) Curcumin I (PubChem CID: 969516), (2) Curcumin II (PubChem CID: 5469424) and (3) Curcumin III (PubChem CID: 5315472) and (A) Casein Kinase-1 (PDB Id: 2IZS), (B) Glycogen Synthase Kinase-3β (PDB Id: 3F88). Ligand is represented as stick and interacting amino acid (labeled) of receptor is represented as disc sphere. Dotted line represents type of interactions: Dark green: Hydrogen bond; Light Green: Van Der Waals interaction; Pink: Pi-Alkyl interaction; White: Carbon hydrogen bond; Red: Acceptor-Acceptor interaction; Orange: Pi-Sulphur interaction.

**Figure 6:**
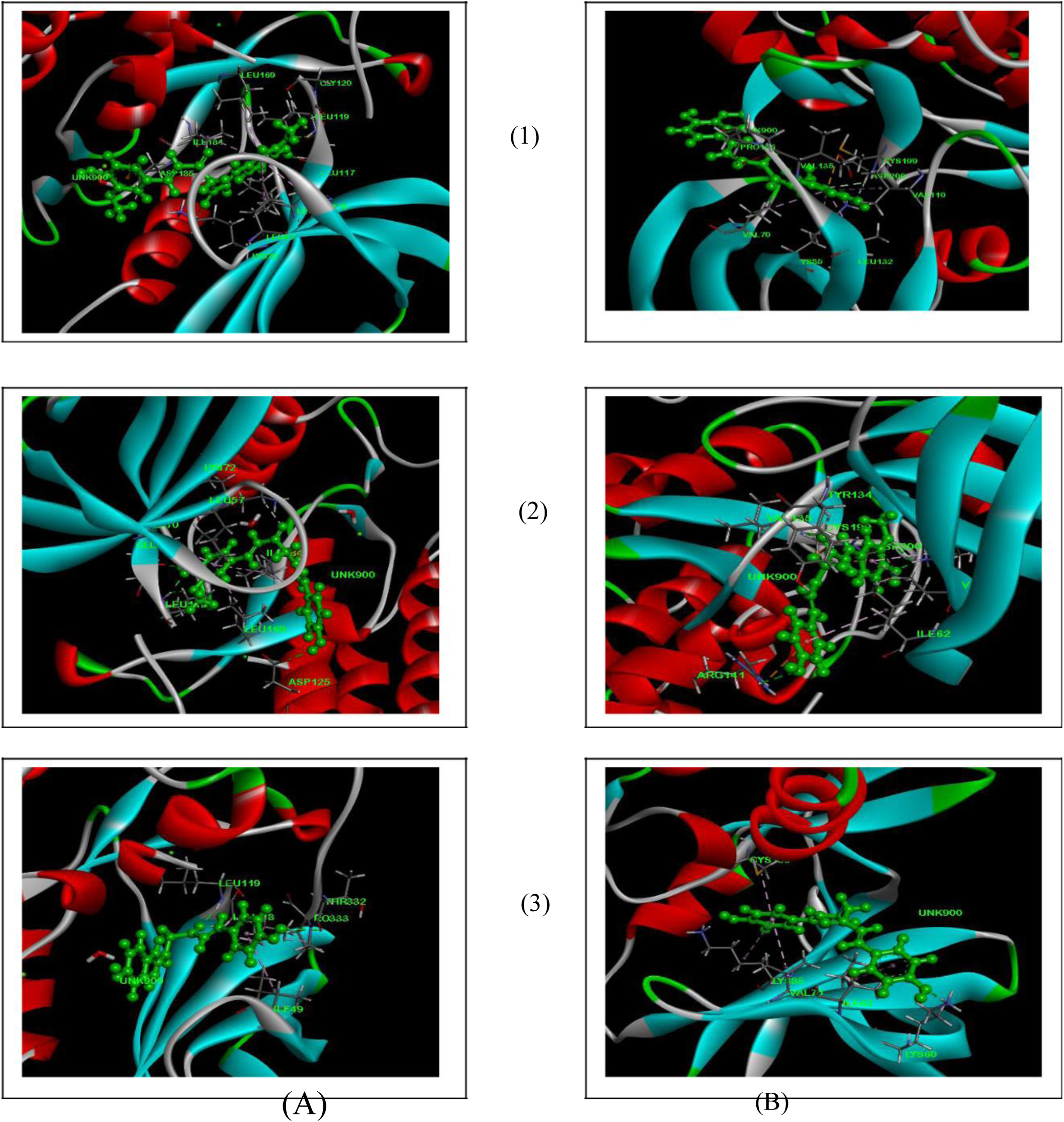
3D representation of best possible poses and ligand-receptor interactions between (1) Curcumin I (PubChem CID: 969516), (2) Curcumin II (PubChem CID: 5469424) and (3) Curcumin III (PubChem CID: 5315472) and (A) Casein Kinase-1 (PDB Id: 2IZS), (B) Glycogen Synthase Kinase-3β (PDB Id: 3F88). Ligand (UNK: 900) is represented as ball and stick and interacting amino acid (labeled) of receptor is represented in stick style. Receptor is represented in ribbon style. Dotted line represents type of interactions: Green: Hydrogen bond; Pink: Pi-Alkyl interaction; White: Carbon hydrogen bond; Red: Acceptor-Acceptor interaction; Orange: Pi-Sulphur interaction.

### 2.2. Structure Based Drug Likeness Property and ADME/Toxicity Prediction

The molecular structures of every ligands were analyzed using SWISSADME server (http://www.swissadme.ch/) in order to confirm whether the physicochemical properties like molecular weight, hydrogen bond donor, hydrogen bond acceptor, molar refractivity etc. of ligands follow Lipinski’s rule of five or not [33]. Additional physicochemical properties such as topological polar surface area, number of rotatable bonds, drug score, drug likeness score etc. of ligand molecules were again calculated using OSIRIS property explorer (https://www.organic-chemistry.org/prog/peo/) [34]. The result of drug likeness property analysis is summarized in **Table 2**. ADME/T profile for each of the ligand molecules was analyzed using an online based server admetSAR (http://lmmd.ecust.edu.cn/admetsar1/predict/) to predict their various pharmacokinetic and pharmacodynamic properties including blood brain barrier permeability, human abdominal adsorption, AMES toxicity, Cytochrome P (CYP) inhibitory promiscuity, carcinogenicity, mutagenicity, Caco-2 permeability etc [35]. The result of ADME/T for all the ligand molecules is represented in **Table 3**.

**Table 2:** Result of drug likeness property analysis of Curcumin I (PubChem CID: 969516), Curcumin II (PubChem CID: 5469424) and Curcumin III (PubChem CID: 5315472). **Lipinski’s rule of five:** Molecular weight: <500, Number of H-bond donors: ≤5; Number of H-bond acceptors: ≤10; Lipophilicity (expressed as LogP): <5; and Molar refractivity: 40-130.

| Drug Likeness Properties | Curcumin I | Curcumin II | Curcumin III |
| --- | --- | --- | --- |
| Molecular Weight | 368.38 g/mol | 338.35 g/mol | 308.33 g/mol |
| LogP | 3.27 | 2.78 | 1.75 |
| LogS | -3.94 | -3.92 | -3.80 |
| H-bond Acceptor | 6 | 5 | 4 |
| H-bond Donor | 2 | 2 | 2 |
| Molar Refractivity | 102.80 | 96.31 | 89.82 |
| Heavy Atoms | 27 | 12 | 23 |
| TPSA | 93.06 Å<sup>2</sup> | 83.83 Å<sup>2</sup> | 74.60 Å<sup>2</sup> |
| Rotatable bonds | 8 | 8 | 6 |
| Drug Likeness Score | -2.63 | -3.55 | -4.48 |
| Drug Score | 0.53 | 0.41 | 0.41 |

**Table 3:**
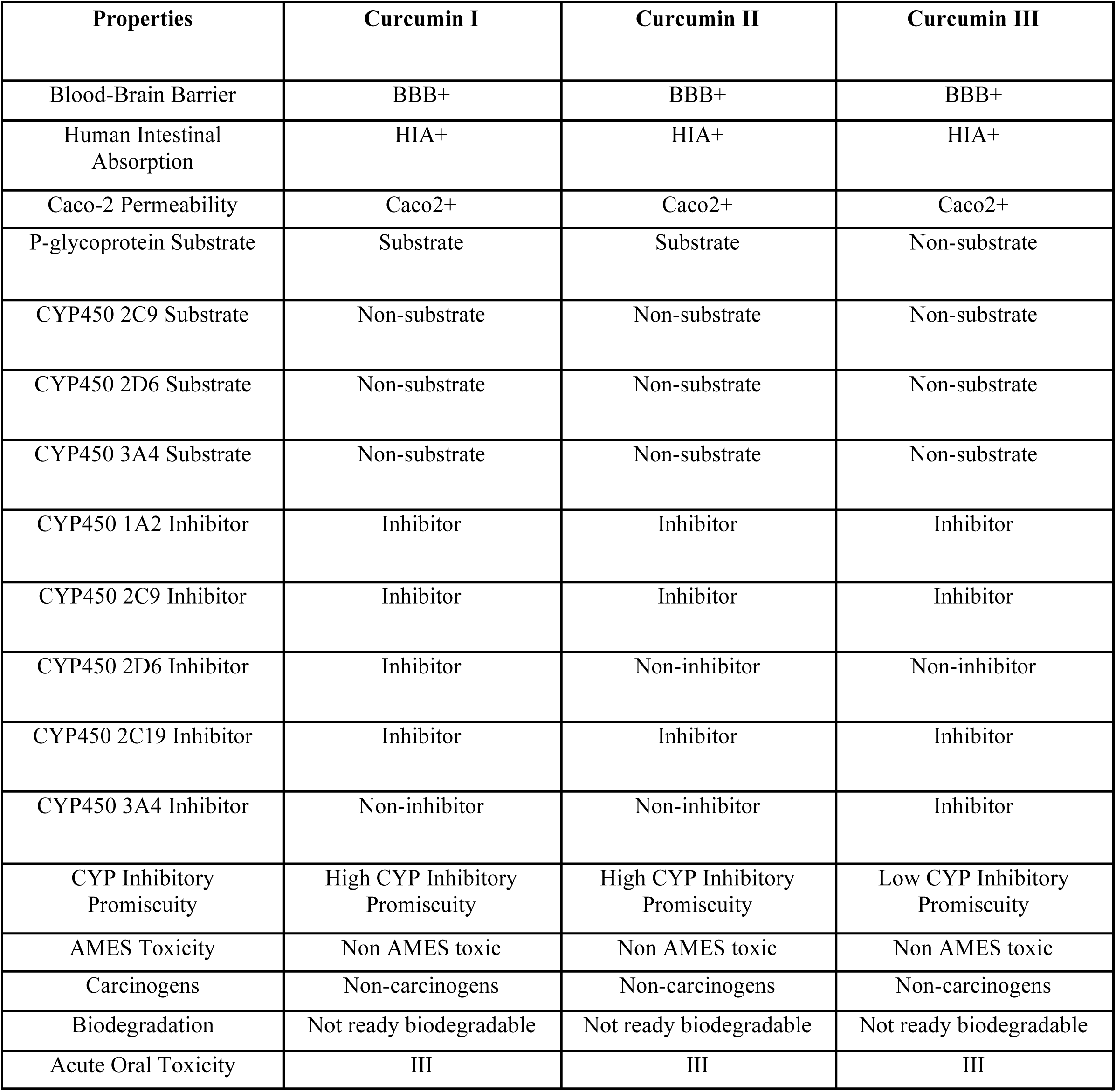
ADME/T properties of Curcumin I (PubChem CID: 969516), Curcumin II (PubChem CID: 5469424) and Curcumin III (PubChem CID: 5315472). BBB+: Capable of penetrating blood brain barrier (BBB); HIA+: Absorbed in human intestinal tissue; Caco-2+: Permeable through the membrane of Caco-2 cell lines; CYP450: Cytochrome P450.

| Properties | Curcumin I | Curcumin II | Curcumin III |
| --- | --- | --- | --- |
| Blood-Brain Barrier | BBB+ | BBB+ | BBB+ |
| Human Intestinal Absorption | HIA+ | HIA+ | HIA+ |
| Caco-2 Permeability | Caco2+ | Caco2+ | Caco2+ |
| P-glycoprotein Substrate | Substrate | Substrate | Non-substrate |
| CYP450 2C9 Substrate | Non-substrate | Non-substrate | Non-substrate |
| CYP450 2D6 Substrate | Non-substrate | Non-substrate | Non-substrate |
| CYP450 3A4 Substrate | Non-substrate | Non-substrate | Non-substrate |
| CYP450 1A2 Inhibitor | Inhibitor | Inhibitor | Inhibitor |
| CYP450 2C9 Inhibitor | Inhibitor | Inhibitor | Inhibitor |
| CYP450 2D6 Inhibitor | Inhibitor | Non-inhibitor | Non-inhibitor |
| CYP450 2C19 Inhibitor | Inhibitor | Inhibitor | Inhibitor |
| CYP450 3A4 Inhibitor | Non-inhibitor | Non-inhibitor | Inhibitor |
| CYP Inhibitory Promiscuity | High CYP Inhibitory Promiscuity | High CYP Inhibitory Promiscuity | Low CYP Inhibitory Promiscuity |
| AMES Toxicity | Non AMES toxic | Non AMES toxic | Non AMES toxic |
| Carcinogens | Non-carcinogens | Non-carcinogens | Non-carcinogens |
| Biodegradation | Not ready biodegradable | Not ready biodegradable | Not ready biodegradable |
| Acute Oral Toxicity | III | III | III |

### 2.6. Structure Based Pharmacological Activity and P450 Site of Metabolism Prediction

Biological activities of each ligand molecule were predicted using PASS (Prediction of Activity Spectra for Substances) online server (http://www.pharmaexpert.ru/passonline/) [36]. All of the ligand molecules were analyzed with respect to probability of activity (Pa) and inactivity (Pi) for 25 intended biological activities depending on their structure. Results of PASS prediction are summarized in **Table 4**. P450 site of metabolism of each ligand molecule was predicted using SMARTcyp online server (https://smartcyp.sund.ku.dk/mol_to_som) in order to analyze which atoms in the molecule are more prone to metabolism by Cytochrome P450 family of enzymes-CYP3A4, CYP2D6, CYP2C9 based on specific ligand structure [37]. Result of Cytochrome P450 site of metabolism prediction is summarized in **Table 5** and **Figure 7**.

**Table 4:** Pharmacological activities of Curcumin I (PubChem CID: 969516), Curcumin II (PubChem CID: 5469424) and Curcumin III (PubChem CID: 5315472). When Pa>0.7: Compound is very likely to exhibit the activity; When 0.7>Pa>0.5: Compound is likely to exhibit the activity; When Pa<0.5: Compound is less likely to exhibit the activity.

| Sl. No. | Biological Activity | Curcumin I |  | Curcumin II |  | Curcumin III |  |
| --- | --- | --- | --- | --- | --- | --- | --- |
|  |  | Pa | Pi | Pa | Pi | Pa | Pi |
| 01. | Antiallergic | 0.435 | 0.043 | 0.408 | 0.050 | 0.251 | 0.129 |
| 02. | Antiinflammatory | 0.677 | 0.019 | 0.667 | 0.020 | 0.704 | 0.015 |
| 03. | Antimutagenic | 0.814 | 0.004 | 0.834 | 0.003 | 0.790 | 0.004 |
| 04. | Apoptosis agonist | 0.803 | 0.008 | 0.861 | 0.005 | 0.871 | 0.005 |
| 05. | Antiulcerative | 0.651 | 0.007 | 0.650 | 0.007 | 0.633 | 0.009 |
| 06. | Antioxidant | 0.610 | 0.004 | 0.624 | 0.004 | 0.637 | 0.004 |
| 07. | Caspase 3 stimulant | 0.747 | 0.009 | 0.719 | 0.010 | 0.492 | 0.029 |
| 08. | Chemopreventive | 0.692 | 0.007 | 0.718 | 0.006 | 0.576 | 0.011 |
| 09. | Carminative | 0.833 | 0.003 | 0.810 | 0.004 | 0.822 | 0.003 |
| 10. | DNA ligase (ATP) inhibitor | 0.314 | 0.040 | 0.326 | 0.036 | 0.364 | 0.024 |
| 11. | Free radical scavenger | 0.766 | 0.003 | 0.785 | 0.003 | 0.670 | 0.004 |
| 12. | Fibrinolytic | 0.731 | 0.013 | 0.735 | 0.012 | 0.717 | 0.017 |
| 13. | Gluconate 2-dehydrogenase inhibitor | 0.833 | 0.010 | 0.822 | 0.012 | 0.836 | 0.009 |
| 14. | Hepatoprotectant | 0.469 | 0.023 | 0.656 | 0.009 | 0.578 | 0.014 |
| 15. | HMOX1 expression enhancer | 0.826 | 0.003 | 0.854 | 0.003 | 0.858 | 0.003 |
| 16. | Membrane permeability inhibitor | 0.724 | 0.029 | 0.736 | 0.025 | 0.721 | 0.030 |
| 17. | Prostate cancer treatment | 0.554 | 0.006 | 0.555 | 0.006 | 0.573 | 0.005 |
| 18. | Proliferative diseases treatment | 0.558 | 0.014 | 0.590 | 0.011 | 0.553 | 0.014 |
| 19. | Reductant | 0.864 | 0.003 | 0.884 | 0.003 | 0.908 | 0.003 |
| 20. | Radioprotector | 0.559 | 0.019 | 0.561 | 0.019 | 0.566 | 0.019 |
| 21. | Sigma receptor agonist | 0.505 | 0.024 | 0.521 | 0.022 | 0.488 | 0.027 |
| 22. | Sugar-phosphatase inhibitor | 0.593 | 0.057 | 0.555 | 0.067 | 0.782 | 0.019 |
| 23. | TNF expression inhibitor | 0.764 | 0.004 | 0.901 | 0.002 | 0.825 | 0.003 |
| 24. | UDP-glucuronosyltransferase substrate | 0.747 | 0.011 | 0.775 | 0.009 | 0.739 | 0.012 |
| 25. | Vasoprotector | 0.678 | 0.012 | 0.620 | 0.017 | 0.654 | 0.014 |

**Table 5:** Result of P450 metabolism site prediction. Atoms are ranked on the basis of lowest enzyme score which means highest probability of being catalyzed by enzyme. *Enzyme Score: Energy - 8*A - 0*.*04*SASA*. A: Accessibility (not shown) is the relative distance of atom from centre of molecule. Energy: approximate activation energy required for CYP active site to catalyze particular atom. SASA: Solvent Accessible Surface Area means local accessibility of atom. 2DSASA: value calculated from molecular topology.

| Compounds | Enzymes | Ranking | Atom | Enzyme Score | Energy | 2DSASA |
| --- | --- | --- | --- | --- | --- | --- |
| Curcumin I | CYP3A4 | 1. | C.13 | 43.6 | 48.5 | 22.4 |
|  |  | 2. | C.26 | 51.6 | 62.2 | 64.3 |
|  |  | 3. | C.23 | 66.5 | 74.1 | 27.8 |
|  | CYP2D6 | 1. | C.1 | 59.6 | 62.2 | 64.3 |
|  |  | 2. | C.13 | 74.5 | 48.5 | 22.4 |
|  |  | 3. | C.22 | 89.5 | 77.2 | 28.6 |
|  | CYP2C9 | 1. | C.1 | 59.6 | 62.2 | 64.3 |
|  |  | 2. | C.13 | 71.3 | 48.5 | 22.4 |
|  |  | 3. | C.22 | 87.9 | 77.2 | 28.6 |
| Curcumin II | CYP3A4 | 1. | C.13 | 43.6 | 48.5 | 22.4 |
|  |  | 2. | C.1 | 51.6 | 62.2 | 64.3 |
|  |  | 3. | C.6 | 66.5 | 74.1 | 27.8 |
|  | CYP2D6 | 1. | C.1 | 59.6 | 62.2 | 64.3 |
|  |  | 2. | C.13 | 74.5 | 48.5 | 22.4 |
|  |  | 3. | C.22 | 89.3 | 77.2 | 31.3 |
|  | CYP2C9 | 1. | C.1 | 59.6 | 62.2 | 64.3 |
|  |  | 2. | C.13 | 71.3 | 48.5 | 22.4 |
|  |  | 3. | C.22 | 87.7 | 77.2 | 31.3 |
| Curcumin III | CYP3A4 | 1. | C.11 | 43.6 | 48.5 | 22.4 |
|  |  | 2. | C.2 | 68.9 | 77.2 | 31.3 |
|  |  | 3. | C.1 | 73.2 | 80.8 | 28.6 |
|  | CYP2D6 | 1. | C.11 | 74.5 | 48.5 | 22.4 |
|  |  | 2. | C.2 | 89.3 | 77.2 | 31.3 |
|  |  | 3. | C.1 | 99.8 | 80.8 |  |
|  | CYP2C9 | 1. | C.11 | 71.3 | 48.5 | 22.4 |
|  |  | 2. | C.2 | 87.7 | 77.2 | 31.3 |
|  |  | 3. | C.1 | 97.4 | 80.8 | 28.6 |

**Figure 7:**
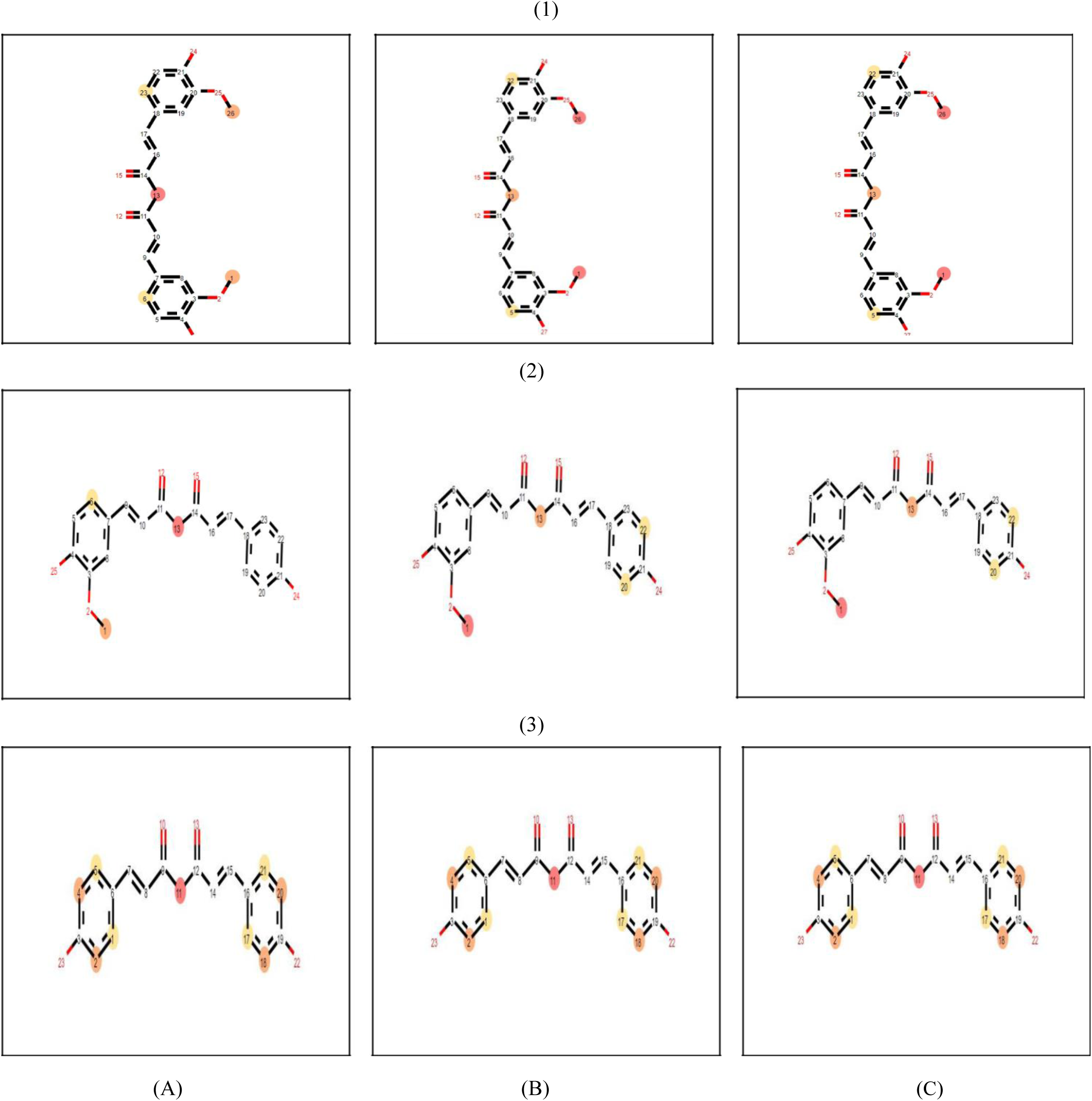
Representation of best possible atoms (marked in colored spheres) of (1) Curcumin I (PubChem CID: 969516), (2) Curcumin II (PubChem CID: 5469424) and (3) Curcumin III (PubChem CID: 5315472) subjected to metabolism by CYP450 group of enzymes (A) CYP 3A4, (B) CYP 2D6 and (C) CYP 2C9. Atoms marked in darker sphere are subjected to have better probability of metabolism by enzymes followed by ones marked in lighter sphere.

## 3. Result

### 3.1. Molecular Docking

All of the selected ligand molecules docked successfully with both CK1 and GSK3B with notable binding energies (**Table1**). Curcumin I docked with a binding energy of −10.31 Kcal/mol within the binding site of CK1. It formed 1 conventional hydrogen bond with Glu117 with 1.62 Å distance apart and 2 conventional hydrogen bonds with Leu119 with 1.99 Å and 2.44 Å distance apart respectively in the binding site backbone of CK1. In addition, Curcumin I also formed two nonconventional hydrogen bonds with Lys72 and Gly140. Moreover, it interacted with 10 amino acid residues in total along with Pi-Alkyl (Hydrophobic) and Pi-Anion (Electrostatic) interactions. On the contrary, Curcumin I interacted with GSK3B with slightly lower energy (−7.533 Kcal/mol) than CK1 within the binding pocket. It formed 2 conventional hydrogen bonds with Lys85 at 2.22 Å and 2.84 Å distance apart and 1 conventional hydrogen bond with Val135 at 2.54 Å distance apart inside the binding site of GSK3B. Curcumin I interacted with 8 amino acid residues within the binding site of GSK3B and notably with other additional types of interactions like-Pi-Sulfur, Pi-Alkyl interactions. It also formed 2 non-conventional hydrogen bonds with Pro136 and Asp200 with GSK3B.

Curcumin II docked to CK1 with a binding energy of −7.826 Kcal/mol. It formed 4 conventional hydrogen bonds-2 with Leu119 at 1.99 Å and 2.79 Å distance apart and 1 with Glu117 and Asp125 each at 1.84 Å and 2.09 Å distance apart respectively within the binding site of CK1. Moreover, it also formed 1 nonconventional hydrogen bond with Lys72 in the binding pocket backbone. Curcumin II interacted with 8 amino acid residues in total with additional hydrophobic (Pi-Alkyl) binding interactions. Conversely, Curcumin II showed a subtle lower binding energy (−7.588 Kcal/mol) with GSK3B. In the binding site backbone of GSK3B it formed 2 conventional hydrogen bonds with Lys85 at 1.87 Å, 2.97 Å distance apart and another conventional hydrogen bond with Val135 at 2.05 Å distance apart. Curcumin II also formed 2 non-conventional hydrogen bonds with Tyr134 and Asp200 within the binding site of GSK3B. With additional Pi-Alkyl, Pi-Cation and Pi-Sulfur bonding interactions Curcumin II interacted with 8 amino acid residues in total.

Curcumin III docked successfully with CK1 with −6.356 Kcal/mol binding energy. It formed 2 conventional hydrogen bonds with Leu119 and Thr332 at 2.00 Å and 2.71 Å distance apart respectively within the binding pocket. It also formed 2 non-conventional hydrogen bonds with Leu118 and Pro333. Curcumin III interacted with 5 amino acids in total with one hydrophobic interaction (Pi-Alkyl) with Pro333. On the other hand, Curcumin III docked with GSK3B with slightly higher binding energy (−6.618 Kcal/mol). It formed just 1 conventional hydrogen bond with Lys60 at 1.93 Å distance apart. In addition, it interacted with total 5 amino acid residues inside the binding pocket of GSK3B only with Pi-Alky hydrophobic interaction. Curcumin III didn’t show any non-conventional hydrogen bonding with any amino acid residues.

### 3.2. Drug Likeness Property

The results of drug likeness property analysis are summarized in Table 2. All of the selected ligand molecules followed Lipinski’s rule of five with respect to molecular weight (acceptable range: <500), number of hydrogen bond donors (acceptable range: ≤5), number of hydrogen bond acceptors (acceptable range: ≤10), lipophilicity (expressed as LogP, acceptable range: <5) and molar refractivity (40-130) [38]. Curcumin III showed lowest Topological Polar Surface Area (TPSA) of 74.60 Å^2^ followed by Curcumin II (83.83 Å^2^) and Curcumin I (93.06 Å^2^). Again Curcumin III showed lowest LogS value (−3.80) among the three ligand molecules. Curcumin II (−3.92) and Curcumin I (−3.934) showed almost similar LogS value. Curcumin I showed the highest drug likeness score (−2.63) followed by Curcumin II (−3.55) and Curcumin III (−4.48). Curcumin II and Curcumin III showed same drug score (0.41) whereas Curcumin I showed highest drug score (0.53) among the the ligand molecules. Curcumin I and II have 8 rotatable bonds and on the contrary, Curcumin III has 6 rotatable bonds. Curcumin I, II and III have 27, 12 and 23 heavy atoms respectively.

### 3.3. ADME/T Test

Results of ADME/T test are summarized in **Table 3**. All the selected ligand molecules showed blood brain barrier and Caco2 cell membrane permeability. All of them showed positive indication of human intestinal absorption. Curcumin I and II are substrates of cell membrane P-glycoprotein whereas Curcumin III is a non-substrate. All of them are non-substrate for CYP450 3A4, CYP450 2C9 and CYP450 2D6 metabolic enzymes. Curcumin II and III are non-inhibitors of CYP450 2D6 whereas Curcumin I is an inhibitor. All of the selected ligand molecules are inhibitors of CYP450 1A2, CYP450 2C9 and CYP450 2C19 metabolic enzymes. All of the selected ligand molecules exhibited high CYP450 inhibitory promiscuity, no AMES toxicity and non-carcinogenicity. All of them are not readily biodegradable and showed type III acute oral toxicity.

### 3.4. Pharmacological Activity Prediction

The results of pharmacological activity prediction are summarized in **Table 4**. Curcumin I, II and III were analyzed for 25 intended biological activities. All selected ligand molecules showed almost similar pharmacological activities although their scores varied slightly with respect to specific activity. Curcumin II showed 13 pharmacological activities with a Pa score greater than 0.7 whereas Curcumin I and III showed 12 pharmacological activities each with a Pa score greater than 0.7.

### 3.5. P450 Site of Metabolism Prediction

The result of P450 metabolism prediction is summarized in **Table 5** and depicted using **Figure**. Carbon13 and Carbon1 were the most prominent atoms in Curcumin I and II which exhibited lowest enzyme scores for metabolism by 3 isoforms (3A4, 2D6 and 2C9) of CYP450 family of enzymes. And in case of Curcumin III, Carbon11 showed the lowermost enzyme scores for all the isoforms of metabolic enzymes. However, Carbon2, Carbon6, Carbon22, Carbon23 and Carbon26also showed satisfactory enzyme score indicating the probability of being catalyzed by all three enzymes.

## 4. Discussion

Medicinal plants are the source of novel phytocompounds which provide numerous therapeutic benefits. Many plant derived compounds and plant extract have been shown to have wound healing activity [39]. *Curcuma longa* has already been reported in the laboratory experiment to exhibit wound healing activity [40] [41]. The involvement of Wnt signaling and upregulation of this signaling pathway in cutaneous wound healing and injury repair process have been demonstrated in laboratory experiment [42][43]. In this study Curcuminoids (**Figure 3**) (Curcumin I, II and III) from *Curcuma longa* were docked against 2 components (**Figure 4**) (Casein Kinase 1 and Glycogen Synthase Kinase 3 Beta) of Wnt/β-catenin signaling pathway based on the hypothesis that the ligand molecules bind to the target and might augment their activity.

Molecular docking is a method of estimating preferred orientation of small molecule when bound to the binding site of a second molecule. It is one of the most frequently used techniques in structure-based drug designing [44]. Molecular docking works on specific scoring algorithm and assigns binding energy to the ligand molecules that fit with the target which reflects the binding affinity. The low binding energy of a ligand molecule with target indicates high stability of the ligand-receptor complex meaning they remain more time in contact [45]. In this experiment Curcumin I interacted with CK1 with lowest binding energy (−10.311 Kcal/mol) and Curcumin II interacted with GSK3B with lowest binding energy (−7.588 Kcal/mol) suggesting the most favorable binding **(Table 1)**. As a consequence Curcumin I interacted with highest number of amino acid residues (10) in the binding site backbone of CK1 and Curcumin II interacted with 8 amino acid residues within the binding site of GSK3B (**Figure 5 and 6**). Curcumin I also showed almost similar binding energy (−7.533 Kcal/mol) as with Curcumin II for interaction with GSK3B. However, Curcumin III showed highest binding energies for both CK1 and GSK3B binding sites. Hydrogen bonding and hydrophobic interactions play key role by strengthening the drug-receptor interaction. All of the selected ligand molecules formed significant number of hydrogen bonds and hydrophobic interactions within the binding site of the target molecules [46].

Drug likeness property evaluation is an important determinant of the successful drug discovery approach. It helps in specifying physicochemical properties of ligand molecules and helps in determining whether a drug should pass the Phase I clinical trial or not. In this experiment ligand molecules were examined according to Lipinski’s rule of five which states that drugs are likely to have poor bioavailability and lower permeation which violates the rules [38] [47]. All of the selected ligand molecules followed Lipinski’s rule of five. Moreover, 10 or fewer rotatable bonds and topological polar surface area equal (TPSA) to or less than 140 Å^2^ are considered to contribute for better oral bioavailability of the candidate drug molecule [48]. All of the selected ligand molecules have TPSA lower than 140 Å^2^ and rotatable bonds lower than 10. Higher molecular weight reduces the permeability of drug through lipid bilayer and lower ones help in the increment of drug permeability. LogP is expressed in the context of lipophilicity and referred as the logarithm of partition coefficient of the candidate molecule in organic and aqueous phase. Lipophilicity affects the absorption of the candidate drug molecule inside the human body. Higher LogP is associated with lower absorption of the drug inside human body and lower value ensures higher rate of absorption of the candidate drug molecule. LogS value influences the solubility of the candidate molecule and the lowest value is always preferred for the drug molecule under investigation in a drug discovery approach. The number of hydrogen bond donors and acceptors outside the acceptable range again influences the ability of a drug molecule to cross bilayer membrane of cell. All of the selected ligand molecules followed Lipinski’s rule of five in this experiment [49] [50]. All selected molecules showed druggable properties within acceptable range (**Table 2**).

Application of *in silico* ADME/T test has gained attention of researchers over the last decade. *In silico* investigation of adsorption, distribution, metabolism and toxicity has enhanced the in-vitro ADME/T test and thus has reduced the time and cost along with increasing the success rate of drug discovery approach [51-58]. Blood brain barrier permeability becomes major concern when drugs target primarily the cells of central nervous system (CNS). Oral delivery system is the most commonly used route of drug administration and the administered drug passes through the digestive tract so it is appreciable that the drug is highly absorbed in human intestinal tissue. P-glycoproteins are embedded on the cell membrane facilitate the transport of many drugs inside the cell and therefore its inhibition may affect the normal drug transport. *In vitro* study of drug permeability test utilizes Caco2 cell line and its permeability to the intended candidate drug molecule reflects that the drug is easily absorbed in the intestine [59]-[61]. All Curcumin analogues were reported to be non-inhibitory to P-glycoprotein, permeable through blood brain barrier, absorbable in the human intestinal tissue and permeable through membrane Caco2 cell lines. Cytochrome P450 family of enzymes plays major role in drug interaction, metabolism and excretion inside the body. Inhibition of these enzymes may lead to the elevation of drug toxicity, slow clearance and malfunction of the drug compound [62]. Inhibitory effect of all three Curcumin analogues was observed against multiple enzymes which may lead to poor degradability and slow excretion of those compounds inside human body. The acute oral toxicity is expressed in terms of median lethal dose (LD_50_), the dose that is capable of killing 50% of the animals under study within 24 hours. Chemicals can be classified into 4 categories (I to IV) based on their extent to induce oral toxicity [63][64]. All selected ligand molecules showed type III oral toxicity. Mutagenicity is considered as one of the most common end point of toxicity. AMES toxicity examines the toxicity of chemicals [65][66]. None of the selected ligand molecules showed AMES toxicity and carcinogenicity (**Table 3**).

Prediction of Activity Spectra for Substances (PASS) predicts the biological activity spectrum of a compound based on its native structure. PASS predicts the activity of a compound based on Structure Activity Relationship Base (SAR Base) which assumes that the activity of a compound is related to its structure. It functions by comparing the 2D structure of a compound relative to another well known compound having biological activities existing in the database with almost 95% accuracy [67]. It predicts the result in the context of probability of activity (Pa) and Probability of inactivity (Pi) of a compound and the result varies between 0.000 and 1.000. When Pa>Pi only then the activity is considered possible for a compound [68]. When Pa>0.7, then the compound is very likely to exhibit the activity but possibility of the compound being analogue to a known pharmaceutical is also high. When 0.5< Pa <0.7, then the compound is likely to exhibit the activity but the probability is less along with the chance of being a known pharmaceutical agent is also lower. When Pa<0.5, then the compound is less likely to exhibit the activity [69]. Curcumin II was reported to be more biologically active than other two compounds for 25 selected activities (**Table 4**).

Cytochrome P450 (CYP450) family comprises 57 isoforms of enzymes responsible for xenobiotic metabolism inside human body. So, rigorous testing of compound’s probability of being metabolized by the CYP450 enzymes is a prerequisite for drug discovery approach. *In silico* approach for CYP450 utilizes 2D structure of a compound to determine which site in a molecule is more liable to metabolism by CYP450 enzymes [70] [71]. Curcumin I showed better result in P450 site of metabolism prediction (**Table 5**) (**Figure 7**).

All the selected ligand molecules docked successfully with both targets and therefore all of them might have roles in wound healing. However, Curcumin I showed highest affinity of binding (−10.311 Kcal/mol) with CK1 and Curcumin II showed highest affinity of binding (−7.588 Kcal/mol) with GSK3B which may indicate their better potentiality to play significant role in wound healing. All of the selected ligand molecules performed well in drug likeness property analysis test and ADME/T test but their possibility of metabolism by CYP450 enzymes was poor which may make their chance questionable to consider as a drug. However, further investigation and intervention might be required to improve their metabolism and excretion profile inside human body. All Curcumin analogues showed some significant biological activities in PASS prediction test which should strengthen their possibility to be considered as drug.

Considering all the parameters of the tests in this experiment it can be concluded that, Curcumin I is the best enhancer of CK1 in and Curcumin II is the best enhancer of GSK3B in Wnt signaling pathway. However, other compounds should also be investigated further since they also performed well in docking experiment. Further *in vitro* and *in vivo* studies are required to confirm the roles of Curcumin analogues in wound healing.

## 5. Conclusion

Three Curcumins from *Curcuma longa* were utilized in this experiment in a search for a drug to be used in wound healing and injury repair process. Several tests indicated positive result suggesting that Curcuma longa could be a great source of herbal drug for wound healing and injury repair along with some other diseases as predicted in PASS. Hopefully, this study will raise research interest among researchers.

## Acknowledgements

Authors are thankful to Swift Integrity Computational Lab, Dhaka, Bangladesh, a virtual platform of young researchers for providing the tools.

